# Productive Mayaro Virus Infection Requires Host Fatty Acid Synthase for nsP1 S-palmitoylation

**DOI:** 10.64898/2026.07.24.740668

**Authors:** Paola N. Loperena González, Adam Brynes, Bryan Perez Soto, Jacqueline Corry, Taanishi Gulati, Hannah Tsui, Nicholas M. Chesarino, Jesse J. Kwiek

## Abstract

To date, twenty-seven pathogenic human viruses require host-catalyzed *de novo* fatty acid biosynthesis for replication. This pathway is driven by fatty acid synthase (FASN), which produces palmitate. Palmitate is a precursor for various functions during viral infection, including lipid droplet formation for assembly, beta-oxidation for ATP generation, and post-translational modification of proteins. Whether Mayaro virus (MAYV), an emerging alphavirus that causes debilitating arthritogenic disease, required FASN for infection was unknown. Using genetic and pharmacological approaches in a human cell line and primary cell model, we found that MAYV requires FASN-dependent palmitate synthesis for virion production. To determine how palmitate contributes to infection, we pharmacologically inhibited pathways downstream of FASN and found that only 2-bromopalmitate (2-BP), a protein palmitoylation inhibitor, led to a 94% reduction in MAYV infection. S-palmitoylation is a post-translational modification in which palmitate is attached to sulfur atoms in cysteine residues. In chikungunya virus, a related alphavirus, FASN-dependent palmitoylation of nonstructural protein 1 (nsP1) is essential for membrane association and replication. Consequently, we hypothesized that MAYV nsP1 is palmitoylated in a FASN-dependent manner. Using an alkyne acetate analog, Alk-4, metabolized by FASN into alkyne palmitate, we observed specific labeling of wild-type nsP1 at conserved cysteine residues (C417-419), but not of a cysteine-to-alanine triple mutant. Treatment with TVB-2640 or 2-BP abrogated Alk-4 labeling of wild-type nsP1 during active infection, reinforcing that MAYV protein palmitoylation is a FASN-dependent process. Our findings reveal a conserved mechanism of FASN-dependent protein palmitoylation in alphaviruses and highlight FASN as a potential anti-viral target.

**Importance:** Mayaro virus (MAYV) is a neglected, mosquito-borne tropical virus that causes debilitating pathologies, such as chronic joint pain that can last from months to years. Currently, MAYV transmissions are endemic in sylvatic and peri-urban regions in Central and South America and the Caribbean. However, MAYV has been detected in urban-adapted mosquitos like *Aedes aegypti* (*Ae. aegypti*) and is a concern for potential global spread. Consequently, investigating the mechanisms of MAYV infection is critical to uncover opportunities for antiviral drug development. In this study, we report that MAYV infection requires host fatty acid synthase (FASN) derived palmitate for palmitoylation of the viral non-structural protein 1 (nsP1). In addition, we report that inhibiting FASN with the clinically advanced small molecule TVB-2640 significantly reduced MAYV infection and nsP1 palmitoylation. This study highlights FASN as an essential host factor for MAYV replication and establishes it as a promising therapeutic target for MAYV and related alphaviruses.

## Introduction

MAYV is a single-stranded positive-sense RNA virus from the *Togaviridae* family and genus *Alphavirus*. It is categorized within the Semliki Forest virus (SFV) antigenic complex, which also includes chikungunya virus (CHIKV). MAYV clinical manifestations are similar to CHIKV and include fever, maculopapular rash, headache, diarrhea and most commonly muscle pain (myalgia) and joint pain (arthralgia) that could persist for months to years (1,2). Geographically, MAYV is endemic to South and Central America (3,4) and the Caribbean (5), and is mainly spread in sylvatic and peri-urban regions by the *Haemagogus* mosquitoes. However, in urbanized regions of Brazil, MAYV has been identified in both patient serum (6–8) and infected *Ae. aegypti* and other urban-adapted mosquito species (9–12). The ability of *Aedes* and other urban mosquito species to become infected and transmit MAYV has also been described in laboratory studies (13–16). The worldwide distribution of these mosquitoes highlight a concern for potential global MAYV spread (17–19). Structurally, like other alphaviruses, the MAYV genome is composed of two open reading frames that encode the non-structural proteins (nsP 1, 2, 3, and 4) and structural proteins (E1, E2, E3, capsid, and TF protein) (1,2). Importantly, the non-structural proteins are responsible for forming the viral replication complex, where the viral RNA capping enzyme, nsP1, anchors the complex to the host membrane for double-stranded RNA (dsRNA) replication (1,2).

All viruses rely on host cell proteins to replicate and assemble into infectious virions. One such protein is fatty acid synthase (FASN), and at least 27 viruses from 15 different families require FASN activity to replicate (20), including herpesviruses (21–23), flaviviruses (24,25), coronaviruses (26,27), and retroviruses (28). FASN is a 273-kDa cytosolic enzyme that uses NADPH to catalyze the condensation of acetyl-CoA and malonyl-CoA to yield the 16-carbon saturated fatty acid palmitate (29). Subsequently, palmitate can be oxidized or elongated to form a diverse pool of cellular fatty acids (for review (30)). We recently reviewed three mechanisms by which palmitate and its fatty acid derivatives are used during the viral replication process, specifically for the formation of lipid droplets that can anchor viral replication complexes or generate energy, ATP production via mitochondrial beta-oxidation, and co-or post-translational protein modification by fatty acylation (20). Despite the importance of these mechanisms for other viruses, the contributions of FASN to MAYV replication have not yet been described. Here we show in both culture adapted and primary human cell models that MAYV replication requires FASN, and mechanistically that MAYV nsP1 undergoes FASN-dependent palmitoylation. These results establish that FASN is a MAYV host dependency factor and a viable candidate for antiviral therapeutic development.

## Results

### Mayaro virus infection requires human FASN activity

To determine if MAYV infection requires FASN enzyme activity, we used both genetic and pharmacological strategies to reduce FASN activity. We transfected 293T cells with either FASN-specific (FASN) or non-targeting (NT) siRNAs for 48h and then infected cells with MAYV at MOI 0.01 for 24h. FASN protein knockdown reduced MAYV infection by 92% (15.2-fold reduction) compared to NT siRNA treatment (Fig. 1A). To further test the importance of FASN in MAYV infection, we infected human HAP-1 cell lines that either expressed (WT) or lacked (KO) endogenous FASN. We first monitored when cytopathic effect formation (CPE; e.g. cell rounding and detachment) developed in HAP-1 cells. We observed MAYV-dependent CPE formation in both WT and KO cells at 48 and 72 hours post-infection (hpi), but not 24 hpi (Fig. S1A). Using these established timepoints, we infected both FASN WT and KO HAP-1 cells with varying levels of MAYV and scored CPE formation to calculate TCID_50_/mL values. The calculated TCID_50/_mL value in FASN KO HAP-1 cells was approximately 78% and 62% lower than in WT HAP-1 cells at 48 hpi and 72 hpi., respectively (Fig. 1B and Fig. S1A), indicating that absence of endogenous FASN reduced MAYV replication and CPE formation.

**Figure 1:**
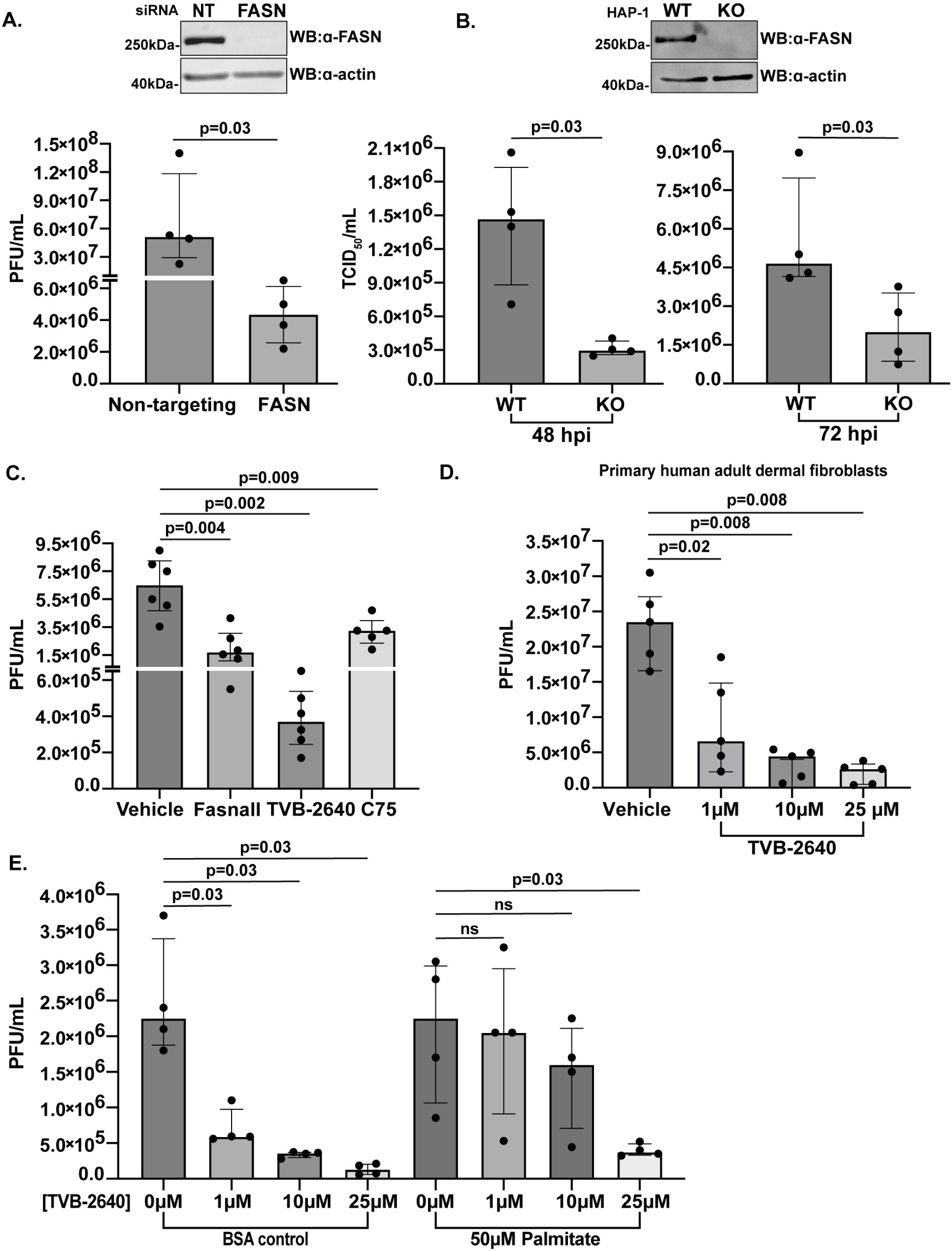
MAYV requires FASN activity for infection. A. 293T cells were transfected with non-targeting (NT) or FASN-specific (FASN) siRNA for 48h and infected with MAYV (MOI 0.01) for 24h. Whole cell protein lysates were immunoblotted for FASN and actin. B. HAP-1 FASN WT or KO cells were infected with MAYV at decreasing MOI for 48h and 72h. CPE was assessed via crystal violet staining and TCID_50_/mL calculated with the Reed-Muench method. Whole cell protein lysates were immunoblotted for FASN and actin. C. 293T cells were pre-treated for 1h with vehicle (DMSO), Fasnall (10 µM), TVB-2640 (25 µM) or C75 (25 µM) for 1hr and then infected with MAYV (MOI 0.01) for 24h. D. Primary human adult dermal fibroblasts (HDFa) were pre-treated for 1h with vehicle (DMSO) or TVB-2640 at 25 µM, 10 µM or 1 µM for 1h and then infected with MAYV (MOI 1) for 24h. E. 293T were pre-treated with DMSO (0 µM) or TVB-2640 at 25 µM, 10 µM or 1 µM in the presence of BSA control or 50 µM of palmitate and then infected with MAYV (MOI 0.01) for 24h. Infectious virus production was assayed by plaque assay in Vero cells in figures A and C to E. Data represented as median with interquartile range of at least four biological replicates. Statistical significance determined by the Mann-Whitney test. Ns = not significant.

Next, we tested the ability of small molecule FASN inhibitors (e.g., Fasnall, TVB-2640, and C75) to block MAYV replication. Fasnall is a thiophenopyrimidine molecule that inhibits HIV-1 infection *in vitro* (28) and demonstrates anti-tumor activity *in vivo* (31). TVB-2640 (denifanstat) is the most clinically advanced FASN inhibitor, currently in clinical trials for metabolic disorders, solid tumors, and acne (32–37). C75 is a FASN inhibitor widely used to investigate the requirements of FASN during viral infection; however, it exhibits known off-target effects (38). We pre-treated 293T cells for 1h with Fasnall (10 µM), TVB-2640 (25 µM), C75 (25 µM), or vehicle control (DMSO) and then infected cells with MAYV at MOI 0.01 for 24h. At concentrations that did not affect cell viability (Fig. S1B-D), we observed that these FASN inhibitors significantly reduced MAYV infection (Fig. 1C). Notably, TVB-2640 resulted in a 94% (16.7-fold) reduction of MAYV infection. To determine if FASN was required for MAYV replication in primary human adult dermal fibroblasts (HDFa), we pre-treated HDFa with TVB-2640 for 1h and infected them with MAYV at MOI 1 for 24h. At all concentrations tested, TVB-2640 inhibited MAYV infection (Fig. 1D), without affecting cell viability (Fig. S1E). Since palmitate is the primary product of FASN activity, we next tested whether supplementation with exogenous palmitate could rescue the TVB-2640-mediated inhibition of MAYV infection. 293T cells were pretreated with TVB-2640 for 1h in the absence or presence of exogenous palmitate, using BSA as the vehicle control, and subsequently infected cells with MAYV at MOI 0.01 for 24h. In the absence of palmitate supplementation, TVB-2640 inhibited MAYV infection in a dose-dependent manner (Fig. 1E). In contrast, when cells were supplemented with 50 µM palmitate, MAYV replication was rescued at both 1 µM and 10 µM of TVB-2640, but not at 25 µM (Fig. 1E). These results demonstrate that FASN-derived palmitate synthesis was required for MAYV infectious virion production.

### MAYV infection does not alter FASN protein expression or localization

Previous studies have shown that viral infection can affect FASN protein levels and in some cases, cause FASN to localize to sites of viral replication (39–42). Among alphaviruses, SFV infection causes FASN to colocalize to type-1 cytopathic vacuoles (CPV-1) (43), and following CHIKV infection, FASN colocalizes with CHIKV dsRNA (44). To determine if FASN protein levels changed during MAYV infection, we infected 293T cells at MOI 1, 5, or 10 for 6h (approximately one viral cycle (45)) or 24h (multiple cycles), and observed that FASN protein levels remained constant during MAYV infection (Fig. 2A). Productive viral infection was confirmed by viral RNA quantification using RT-qPCR (Fig. 2B). Next, to determine if FASN protein localized to sites of MAYV replication, we infected 293T cells with MAYV at MOI 1 for 24h and assessed the subcellular localization of FASN and dsRNA, a MAYV replication intermediate. FASN did not colocalize with dsRNA, as observed by clear distinction of FASN (green) and dsRNA (red) fluorescence (Fig. 2C). Thus, MAYV infection did not change intracellular FASN protein levels or localization.

**Figure 2:**
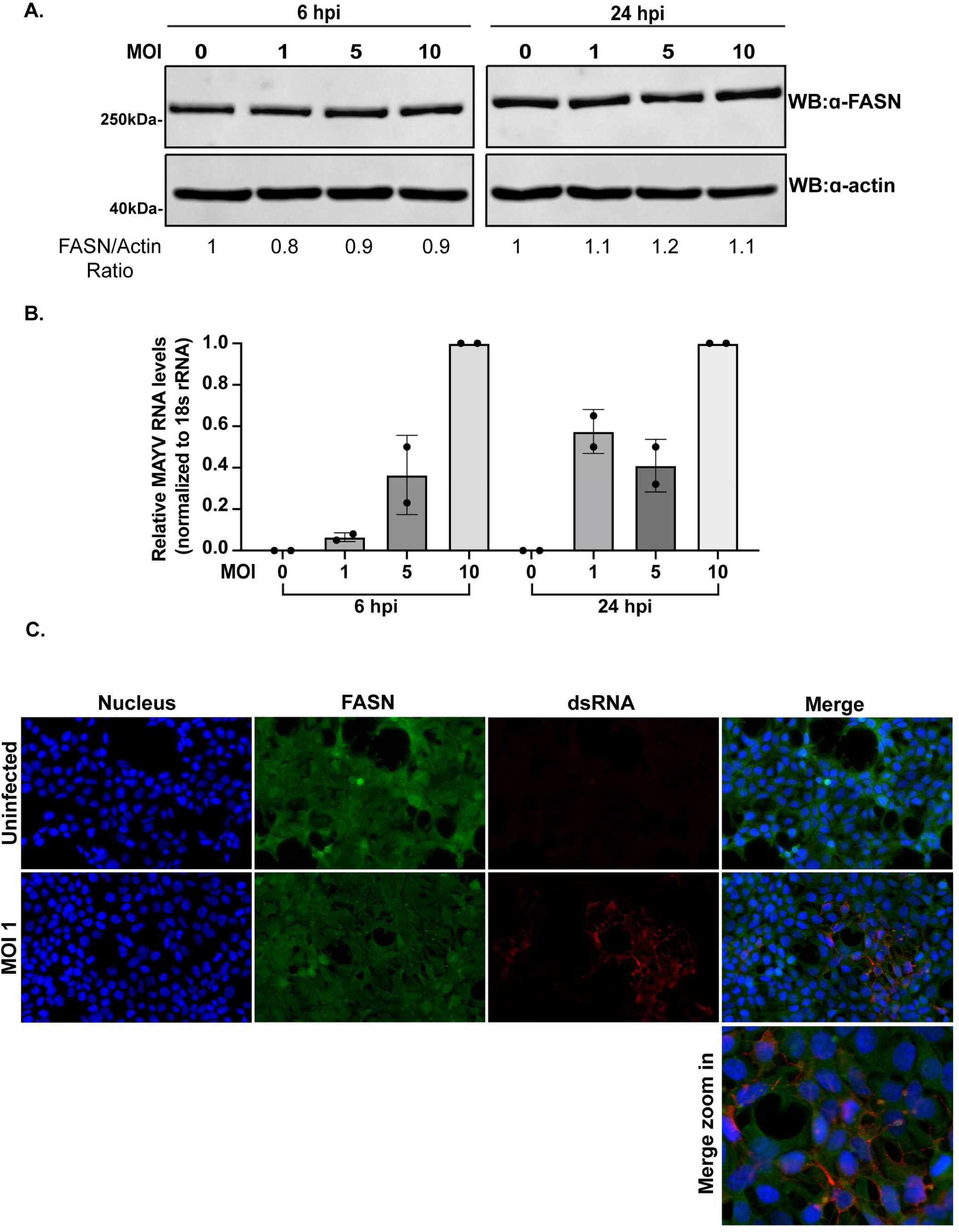
MAYV infection does not affect FASN protein expression and localization. 293T cells were infected with MAYV at MOI 1, 5, or 10 or remined uninfected (MOI 0) for 6h or 24h. Cells were lysed for whole cell protein and RNA collection. A. Protein lysates were immunoblotted for FASN and actin. Densitometry quantification represented as average of two biological replicates. B. The presence of MAYV RNA was assessed by RT-qPCR. Data represented as mean with standard deviation. C. 293T cells were infected with MAYV (MOI 1) for 24h. Cells were fixed and immunolabeled with FASN antibody (green) and J2 (dsRNA) antibody (red). DAPI (blue) marks the nucleus. Images are representative of two biological replicates.

### MAYV nsP1 palmitoylation is a conserved requirement during alphavirus infection

FASN-derived palmitate could contribute to viral replication in at least three ways, such as lipid droplet formation, mitochondrial beta-oxidation, and viral protein acylation. To test if inhibition of these pathways affects MAYV infection, we pre-treated 293T cells for 1h with an inhibitor of lipid droplet formation (A922500,10 µM), an inhibitor of mitochondrial beta-oxidation (etomoxir, 10 µM), or an inhibitor of protein S-palmitoylation (2-bromopalmitate, 2-BP, 12.5 µM) and then infected cells with MAYV at MOI 0.01 for 24h. The inhibitor concentrations used did not affect cell viability (Fig. S2A-C). We observed that MAYV infection was unchanged by A922500 and etomoxir treatment; however, 2-BP treatment reduced MAYV infection by 96% (19.4-fold reduction, Fig. 3A), suggesting a role for protein palmitoylation. Alphaviruses like CHIKV, SFV, and Sindbis virus (SINV) require protein palmitoylation of E glycoproteins, nsP1, and TF (46–51). In CHIKV, nsP1 palmitoylation occurs in a FASN-dependent manner (51). Given the well-characterized role of nsP1 in the alphavirus life cycle (49,52–56), we decided to focus on nsP1 as a target for MAYV protein S-palmitoylation. An amino acid sequence alignment of CHIKV and MAYV nsP1 showed a 70% amino acid identity, including conservation of the CHIKV cysteine palmitoylation sites (C417, C418, C419) (Fig. 3B). Next, we used AlphaFold3 to compare the predicted location of the S-palmitoylation sites between CHIKV and MAYV nsP1 (Fig. 3C). The root mean square deviation (RMSD) value of 0.59Å suggested that both CHIKV and MAYV nsP1 share highly similar predicted conformations. Thus, we hypothesized that MAYV nsP1 would exhibit a similar palmitoylation pattern to CHIKV nsP1. We then evaluated the palmitoylation capacity of MAYV nsP1 using an ectopic expression system. We generated N-terminal Flag-tagged MAYV nsP1 expression vectors (Flag-nsP1) for both the wild-type (WT) protein and a palmitoylation-deficient mutant (C417-419A), in which the cysteines at positions 417, 418, and 419 were mutated to alanine. To determine if MAYV Flag-nsP1 is palmitoylated, we employed copper(I)-catalyzed azide-alkyne cycloaddition (i.e. click chemistry) where an alkyne group maintains the hydrophobic character of the fatty acid while allowing a targeted reaction with an azide-functionalized fluorescent tag (e.g. 5-TAMRA azide) (57–59). We incubated Flag-nsP1-transfected cells with Alk-16, an alkyne palmitate analog, and observed that Alk-16 specifically labeled WT Flag-nsP1 but not the C417-419A mutant (Fig. 3D). Furthermore, treatment with 2-BP (12.5 µM) markedly reduced Alk-16 labeling of WT Flag-nsP1. Initially we observed a lower recovery of the Flag-nsP1 C417-419A following immunoprecipitation. To test that the absence of a palmitoylation signal was not due to lower protein abundance, we transfected a range of plasmid concentrations for each Flag-nsP1 construct. We found that even at higher protein expression levels, Flag-nsP1 C417-419A remained unlabeled (Fig. S2D). Next, to determine if palmitoylation occurs at a specific cysteine within the triad, we generated single and double cysteine-to-alanine mutations within the Flag-nsP1 constructs. Following Alk-16 labeling, we observed that all single-cysteine variants were palmitoylated to similar levels. However, the C417A,C418A double mutant had enhanced labeling compared to other double mutants (Fig. 3E), suggesting a potential preference for C419. Overall, because all single residue substitutions still permitted palmitoylation, we concluded that MAYV nsP1 could be palmitoylated at any of the cysteines in positions 417-419, without an exclusive preference for a single site.

**Figure 3:**
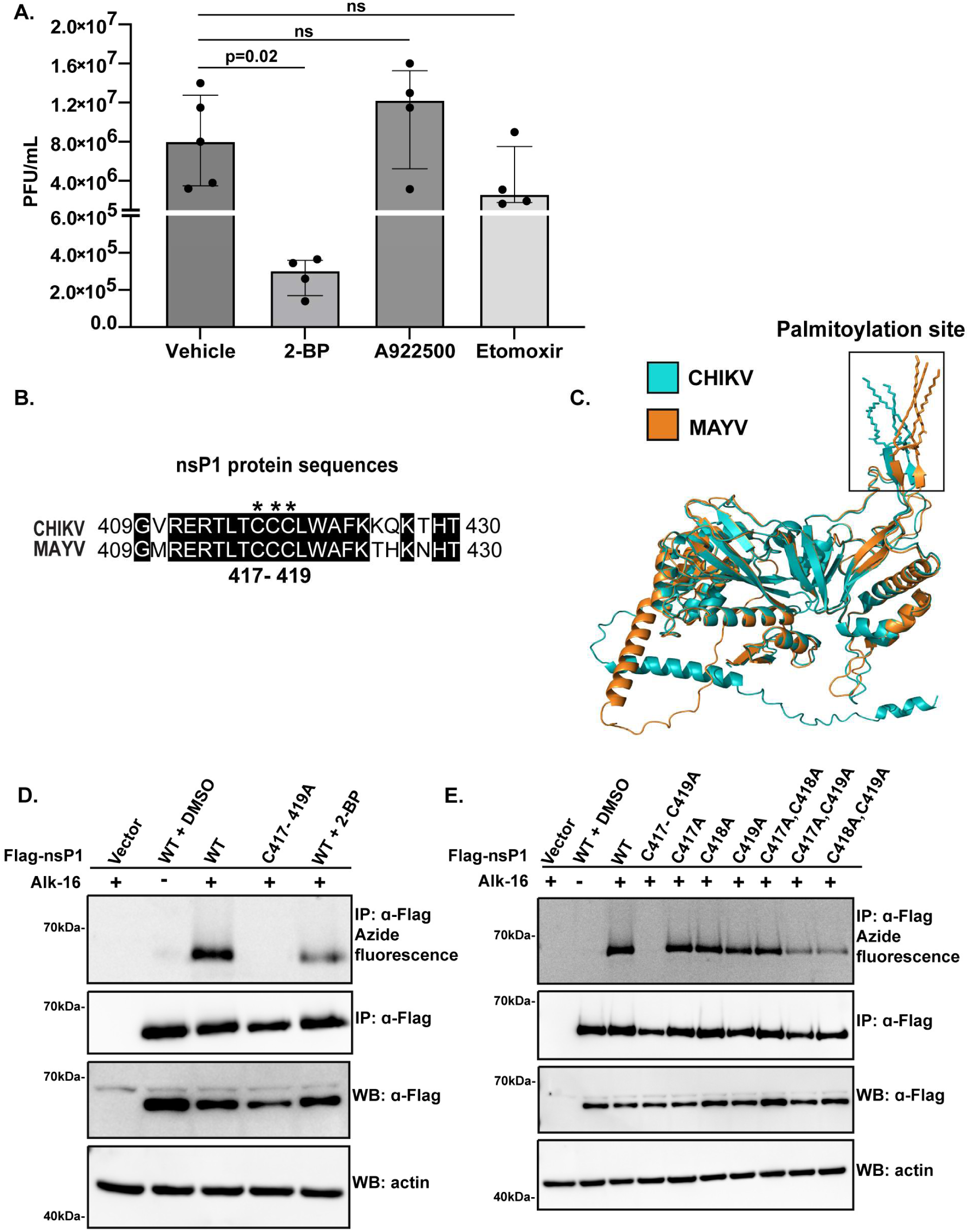
MAYV nsP1 requires protein palmitoylation in cysteines 417 to 419. A. 293T cells were pre-treated with vehicle (DMSO), 2-bromopalmitate (2-BP, 12.5 µM), A922500 (10 µM) and etomoxir (10 µM) for 1h and then infected with MAYV (MOI 0.01) for 24h. Infectious virus production was assayed by plaque assay. Data represented as median with interquartile range of four biological replicates. ns= not significant. B. MAYV nsP1 sequence was compared to CHIKV nsP1 and represented in a protein sequence alignment. Protein alignment generated with Expasy. C. AlphaFold3 prediction of MAYV and CHIKV nsP1 structure including the putative S-palmitoylation site at cysteines 417-419. Predictions were observed and aligned using PyMoL. D. 293T cells were transfected with vector, Flag-nsP1 WT or C417-419A plasmids overnight. Then, 2-BP (12.5 µM) or vehicle (DMSO) control treatment was administered for 6h. After, cells were exposed to DMSO or Alk-16 (50 µM) in the presence of 2% charcoal striped FBS (CS-FBS) for 24h. E. 293T cells were transfected with vector, Flag-nsP1 WT, C417-419A, single or double cysteine mutant plasmids overnight. Cells were exposed to vehicle DMSO or Alk-16 (50 µM) in the presence of 2% CS-FBS for 24h. For figures D and E, after labeling, cells were lysed, Flag-nsP1 was immunoprecipitated and blotted with Flag antibody, and palmitoylation was assessed via click chemistry using 5-TAMRA azide. Whole cell protein lysates were immunoblotted with Flag and actin antibodies. Western blots are representative of two biological replicates.

### MAYV nsP1 undergoes FASN-dependent palmitoylation

Because palmitate is a downstream product of FASN, the palmitate analog Alk-16 does not directly test FASN-dependent protein palmitoylation. Given the critical requirement of FASN during MAYV infection, we investigated whether FASN-derived fatty acids could associate with MAYV nsP1. To study this, we used alkynyl acetic acid (Alk-4), a cell-permeable and click-chemistry-compatible acetate analog that is metabolized by FASN into fatty acids (59). We labeled Flag-nsP1-transfected cells, expressing either WT or C417-419A, with 5 mM or 10 mM Alk-4 and observed labeling on WT Flag-nsP1 but not on the C417-419A mutant (Fig. 4A). In addition, treatment with either the FASN inhibitor TVB-2640 (25 µM) or the palmitoylation inhibitor 2-BP (12.5 µM) markedly reduced Alk-4-dependent nsP1 labeling (Fig. 4B). Together, these results demonstrate that MAYV nsP1 is palmitoylated at cysteines 417-419 in a FASN-dependent manner.

**Figure 4:**
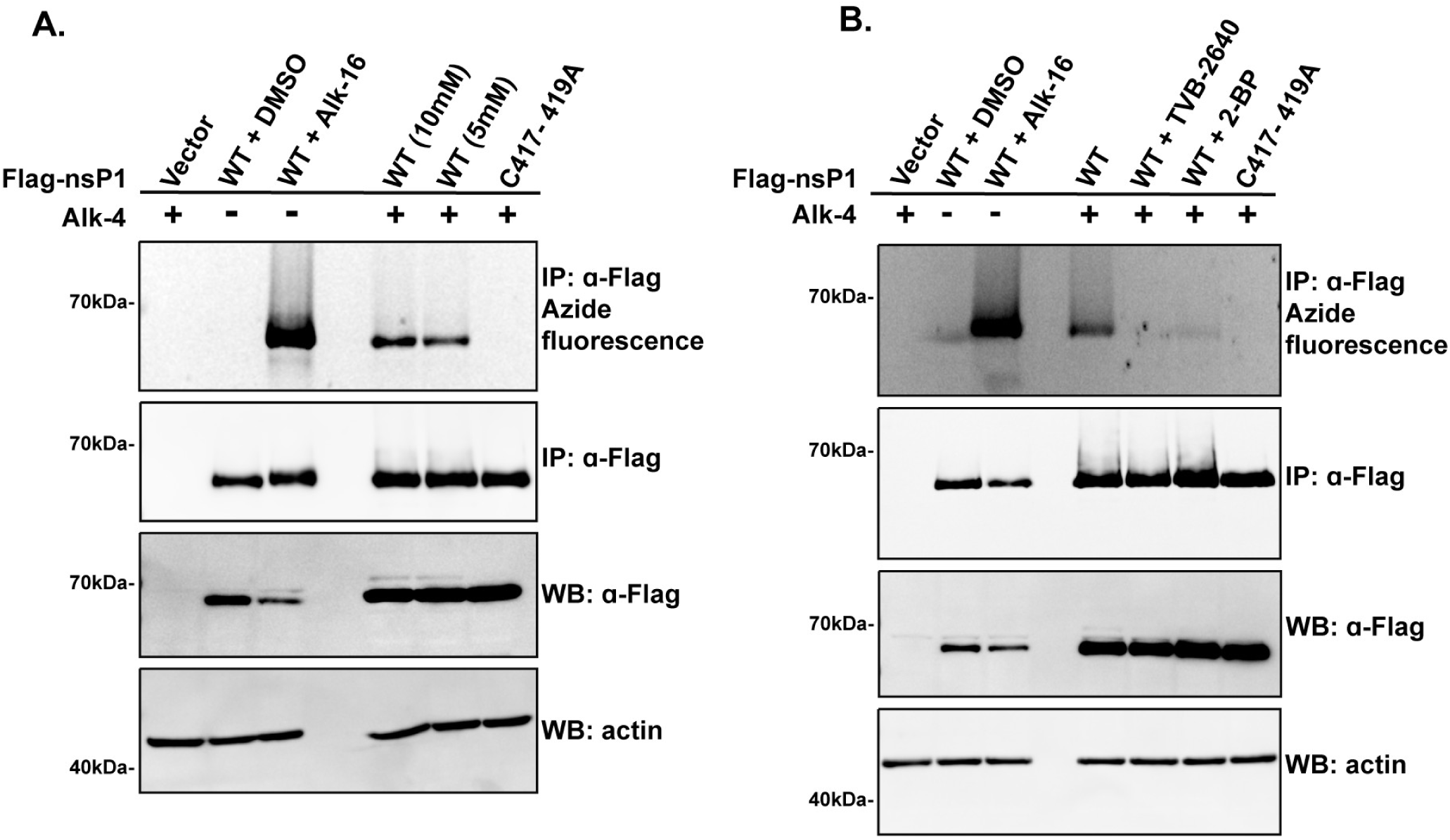
Palmitoylation of ectopic MAYV nsP1 expression is dependent on FASN activity. A. 293T cells were transfected with Flag-nsP1 WT or C417-419A plasmids overnight. Cells were exposed to vehicle (DMSO), Alk-16 (50 µM), or Alk-4 (10 mM or 5 mM) in the presence of 2% charcoal stripped (CS)-FBS for 24h. B. 293T cells were transfected with Flag-nsP1 WT or C417-419A plasmids overnight. Then, TVB-2640 (25 µM), 2-BP (12.5 µM), or DMSO treatment was administered for 6h. After, cells were exposed to DMSO, Alk-16 (50 µM), or Alk-4 (10 mM) in the presence of 2% CS-FBS for 24h. For figures A and B, after labeling, cells were lysed, Flag-nsP1 was immunoprecipitated and blotted with Flag antibody and palmitoylation was assed via click chemistry using 5-TAMRA azide. Whole cell protein lysates were immunoblotted with Flag and actin antibodies. Western blots are representative of two or four biological replicates, respectively.

### FASN activity is required for nsP1 palmitoylation during active MAYV infection

Having established in an ectopic expression system that FASN activity is associated with MAYV nsP1 palmitoylation, we next investigated whether FASN contributed to nsP1 palmitoylation in the context of the full MAYV genome. We modified a previously developed MAYV infectious molecular clone (IMC) (TRVL 4675) (60) to contain an N-terminal Flag-tagged nsP1 (Flag-nsP1 WT IMC) or a corresponding palmitoylation-deficient mutant featuring C417,418,419A mutations (Flag-nsP1 C417-419A IMC) (Fig. 5A). Following transfection into 293T cells, we confirmed comparable Flag expression for both constructs at 72 hours post-transfection (hpt) (Fig. 5B). To assess the functional requirement of nsP1 palmitoylation, we quantified the production of infectious virus 72 hpt. The C417-419A IMC failed to produce infectious virions, as evidenced by the absence of plaques (Fig. 5C and D). Consistent with this observation, robust viral RNA replication was detected via dsRNA staining only in cells transfected with the Flag-nsP1 WT IMC (Fig. S2E). Finally, to test whether MAYV nsP1 palmitoylation is FASN dependent within the context of the full MAYV genome, transfected cells were subjected to Alk-4 labeling in the presence or absence of TVB-2640 or 2-BP. Cells transfected with WT Flag-nsP1 IMC were also independently labeled with Alk-16. Both Alk-16 and Alk-4 labeled Flag-nsP1 encoded by the WT Flag-nsP1 IMC, and, importantly, the Alk-4 signal was reduced by both the FASN inhibitor TVB-2640 (25 µM) and the palmitoylation inhibitor 2-BP (12.5 µM) (Fig. 5E). Due to the severe replication deficiency caused by the cysteine-to-alanine mutations, the C417-419A IMC did not yield sufficient Flag-nsP1 protein to accurately assess its palmitoylation state. Ultimately, these findings demonstrate that MAYV nsP1 undergoes FASN-dependent palmitoylation within a functionally active MAYV replication system.

**Figure 5:**
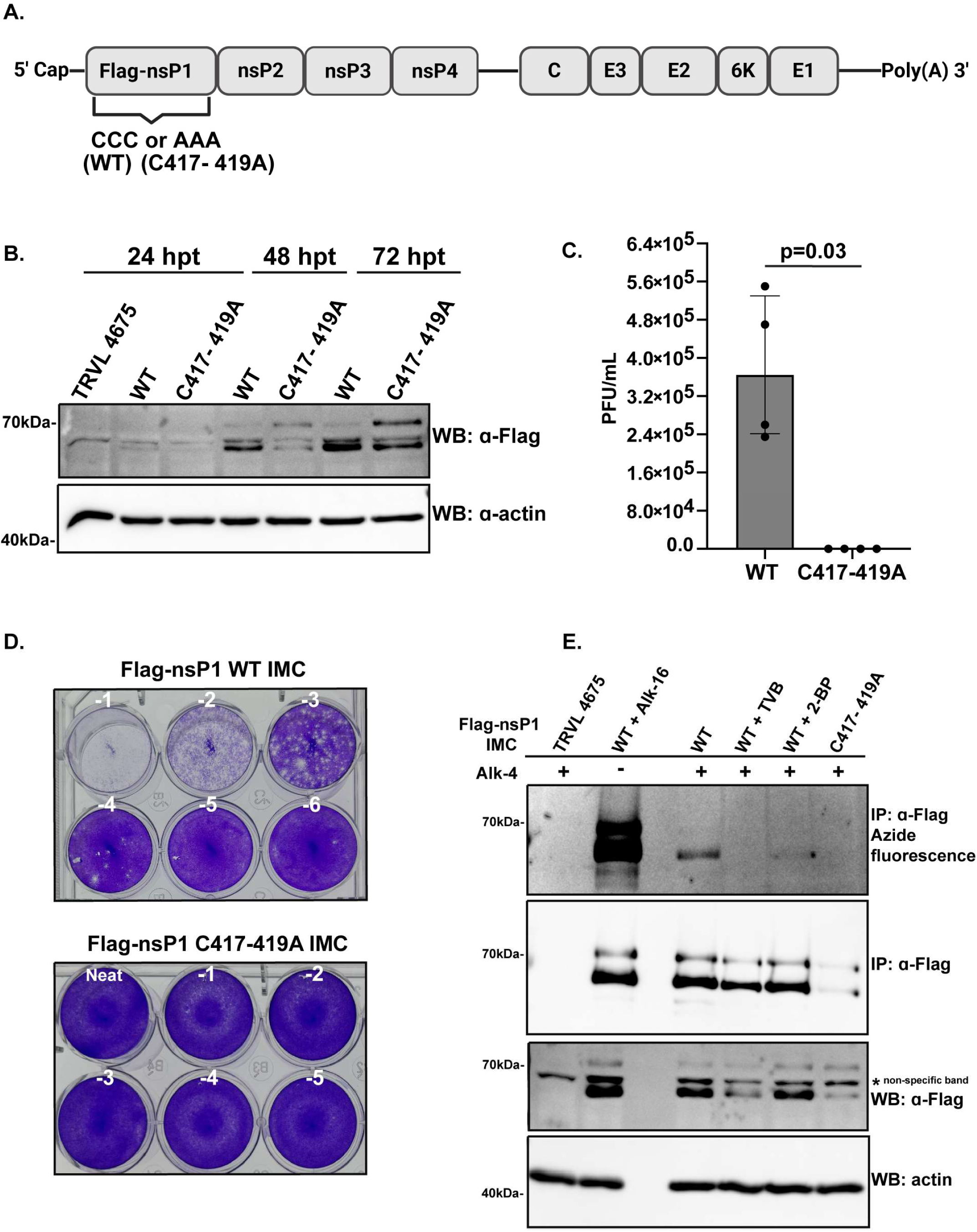
MAYV nsP1 palmitoylation occurs in a FASN-dependent manner during productive infection. A. Schematic representation of the MAYV infectious molecular clone (IMC) constructs with Flag-tagged nsP1 at positions 417, 418, 419 retained as cysteines (WT) or mutated to alanine (C417-419A). Created in BioRender. B. 293T cells were transfected with IMC backbone TRVL 4675 (24h only), Flag-nsP1 WT or C417-419A IMC for 24h, 48h, or 72h. Cells were lysed at each timepoint, and protein lysates were immunoblotted with Flag and actin antibodies. C. 293T cells were transfected with Flag-nsP1 WT or C417-419A IMC for 72h and infectious virus production was assessed by plaque assay. Data represented a median with interquartile range of four biological replicates. Statistical significance determined by the Mann-Whitney test. D. Representative images from plaque assays of Flag-nsP1 WT and C417-419A IMC generated on C. E. 293T cells were transfected with the IMC backbone TRVL 4675, Flag-nsP1 WT or C417-419A plasmids for 48h. Then, Flag-nsP1 WT transfected cells were exposed to TVB-2640 (25 µM) or 2-BP (12.5 µM) treatment for 6h. After, cells were exposed to Alk-16 (50 µM), or Alk-4 (10 mM) in the presence of 2% CS-FBS for 24h. After labeling, cells were lysed, Flag-nsP1 was immunoprecipitated and blotted with Flag antibody, and palmitoylation was assed via click chemistry using 5-TAMRA azide. Whole cell protein lysates were immunoblotted with Flag and actin antibodies. Western blots are representative images of at least three biological replicates.

## Discussion

Here, through a combination of genetic and pharmacological inhibition in both a cell line and a primary cell model, we demonstrate that MAYV replication depends on FASN-derived palmitate synthesis for viral protein palmitoylation. Specifically, treatment with the FASN inhibitor TVB-2640, currently in phase 3 clinical trials, reduced MAYV infection in primary human dermal fibroblasts (Fig. 1D) and abrogated FASN-dependent MAYV nsP1 palmitoylation during active infection (Fig. 5E). These results identify FASN as a MAYV host dependency factor and suggest FASN as a potential alphavirus therapeutic target.

These findings build on earlier reports that used the early-generation FASN inhibitor cerulenin and the lipase inhibitor orlistat to link host fatty acid synthesis to MAYV infection (61). In this study, we evaluated the first-generation FASN inhibitor C75 and second-generation inhibitors Fasnall and TVB-2640 and demonstrated consistent pharmacological reduction of MAYV infection. These findings were complemented by genetic FASN perturbations, wherein both transient FASN knockdown cells and stable FASN knockout cell lines restricted MAYV replication, confirming that the observed antiviral effects are FASN-specific. Finally, the rescue of MAYV replication with exogenous palmitate supplementation establishes that FASN-mediated palmitate synthesis is required for efficient MAYV replication.

In the viral infection cycle, palmitate can support replication by contributing to lipid droplet formation for viral assembly, catabolism by mitochondrial beta-oxidation to generate energy, or by serving as a substrate for post-translational protein fatty acylation (20). Here, we demonstrate that MAYV infection specifically requires protein palmitoylation. These results align with previous studies showing that the alphavirus structural proteins E and TF, as well as the nonstructural protein nsP1, undergo palmitoylation in SINV, SFV and CHIKV (46–51). Furthermore, CHIKV nsP1 palmitoylation has been shown to occur in a FASN-dependent manner (51). Consistent with these findings, our click chemistry assays with Alk-4 demonstrate that during both ectopic nsP1 expression and active MAYV infection, MAYV nsP1 is selectively labeled at cysteines 417, 418, and 419 in a FASN-dependent manner. Although this study focused on nsP1, the sensitivity of MAYV replication to palmitoylation inhibition suggests that other MAYV proteins, such as TF and E, may also undergo FASN-dependent palmitoylation.

During active infection, we observed that nsP1 palmitoylation deficiency abrogated dsRNA synthesis and infectious virion production. This lethal phenotype has also been described for CHIKV (51,62). In contrast, nsP1 palmitoylation deficiency in SINV and SFV does not affect viral particle production (47,63,64) and only attenuates SFV pathogenicity *in vivo* (47). These divergent phenotypes may be explained by the multifunctional roles of nsP1 in alphavirus replication. Aside from functioning as the viral mRNA capping enzyme, nsP1 drives viral replication complex assembly through membrane anchoring (49,52–56). The membrane-binding capacity of alphavirus nsP1 is mediated by both the palmitoylation of a conserved cysteine triad and an amphipathic helix containing a critical tryptophan residue at position 258 (W258) (49). While the amphipathic helix mediates phospholipid membrane interactions, the palmitoylated cysteine triad targets and anchors nsP1 to cholesterol-rich lipid microdomains (49). Although both membrane-binding motifs are conserved across CHIKV, MAYV, SFV, and SINV, membrane binding preferences differ between specific alphaviruses. In SFV and SINV, disrupting palmitoylation reduces nsP1 association with cholesterol-rich lipid microdomains but does not affect virion production (47,56,65,66), whereas mutation of W258 prevents membrane association (49) and abrogates infection (67). Conversely, for CHIKV, mutating only the cysteine triad is sufficient to disrupt lipid microdomain association and completely prevent virion production (51,62). The similar replication-deficient phenotypes observed between MAYV and CHIKV cysteine-to-alanine mutants strongly suggest that FASN-dependent palmitoylation is required to anchor MAYV nsP1 to cholesterol-rich lipid microdomains, thereby driving replication complex formation and dsRNA synthesis. Collectively, our findings highlight a differential dependency on host FASN activity across alphaviruses, even among viruses within the same antigenic complex.

We established that MAYV relies specifically on protein fatty acylation for efficient infection. In contrast, a previous study reported that CHIKV replication in Vero cells was inhibited by the lipid droplet formation inhibitor A922500, the beta-oxidation inhibitor etomoxir, and high concentrations of TVB-2640(68). In our study, we evaluated the contributions of these pathways using lower inhibitor concentrations in human cells, which could account for the phenotypic differences between these studies. We also demonstrated that TVB-2640 maintains antiviral activity at low micromolar concentrations in both a human cell line and primary human cells. These discrepancies underscore how antiviral efficacy can be influenced by drug dosing concentrations, cell type, and species-specific variations.

FASN protein abundance and subcellular localization can be modulated during viral infection (39–42). Here, we found that FASN protein levels remain unchanged during MAYV infection, which has not been previously reported. Additionally, FASN did not localize to sites of dsRNA replication. These data differ from previous findings in SFV and CHIKV infections, where FASN was enriched in SFV replication organelle fractions in infected HeLa cells (43) and FASN co-localized with CHIKV dsRNA in viral replicon particle-infected HeLa cells (44). A study identified that HeLa cells have high basal FASN expression, and FASN upregulation in these cells causes cholesterol reprogramming and subsequent lipid raft remodeling at the plasma membrane (69). Because alphavirus genome replication, particularly for CHIKV, preferentially occurs within cholesterol-rich microdomains in the plasma membrane (49,70), the phenotypic differences observed here may be attributed to cell-type specific variations. Alternatively, these discrepancies might involve the non-canonical role of FASN as an RNA binding protein. During CHIKV infection, FASN directly binds viral RNA; while its enzymatic activity regulates CHIKV RNA levels, FASN can non-enzymatically enhance viral protein translation (71). It has been proposed that CHIKV RNA hijacks FASN to localize its enzymatic activity to viral replication sites, locally elevating palmitate levels to facilitate viral packaging and replication. Whether this mechanism extends to MAYV and other alphaviruses remains unknown.

Our discovery that TVB-2640 blocks FASN-dependent nsP1 palmitoylation and restricts viral infection in primary human cells strengthens the clinical viability of FASN as an antiviral target. TVB-2640, clinically known as denifanstat, has demonstrated a favorable safety and efficacy profile across diverse pathologies, including Phase 3 clinical trials for acne treatment and Phase 2 trials for metabolic dysfunction associated steatohepatitis (MASH) and solid tumors (32–37). In our study, a 1 µM concentration of TVB-2640 was sufficient to restrict MAYV infection in primary human dermal fibroblasts. Clinically, MASH patients or obese patients at risk of MASH treated with TVB-2640 safely achieve drug plasma concentrations of 1-2 µM (33,34). This alignment strongly suggests that the effective antiviral concentrations observed *in vitro* are clinically achievable and well-tolerated in humans.

Currently, MAYV is classified as a neglected tropical mosquito-borne virus with transmission confined to forested and peri-urban regions. However, MAYV has been identified in urban-adapted vectors, including *Ae. aegypti* (9–12). FASN dependency during MAYV infection remains conserved across the transmission cycle, as a previous study showed that MAYV-infected *Ae. aegypti* females have significantly reduced viral titers when FASN is knocked down (72). Because CHIKV and other mosquito-borne infections are endemic in regions where MAYV is present, underdiagnosis and underreporting of MAYV infections are highly probable (73). As

MAYV circulation expands into urban areas, the potential for epidemics increases, underscoring the importance of identifying efficacious antivirals to mitigate future outbreaks. The role of FASN as a host dependency factor makes it a promising antiviral target, as strategies directed against host proteins, rather than viral proteins, may circumvent the rapid evolutionary adaptation that drives conventional drug resistance. In summary, our findings demonstrate that MAYV infection requires FASN activity for nsP1 S-palmitoylation, highlighting FASN as a MAYV host dependency factor and a clinically viable target for anti-alphavirus therapeutic development.

## Acknowledgments

P. N. Loperena González was supported by NIH T32 GM141955. Ideas presented here were conceived and developed while J.J.K. was funded by NIH awards AI090644 and AI141037. We thank Dr. Jacob Yount (OSU) for technical assistance with click chemistry protocols. We thank Dr. James Weger-Lucarelli from Virginia Tech, Blacksburg, VA for providing MAYV infectious molecular clones TRVL 4675 and TRVL 15537. The authors declare no conflict of interest.

## Author Contributions

PNLG: Conceptualization, Methodology, Investigation, Formal analysis, Visualization, Writing - original draft, Writing - review & editing. AB: Microscopy-optimization & figure generation and Writing - review & editing. BPS: computational- AlphaFold3 and sequence alignment figure generation and Writing - review & editing. JC: Viral stock generation and Writing - review & editing. TG: Microscopy-optimization and Writing - review & editing. HT: Cell viability-optimization and Writing - review & editing. NMC: Conceptualization, Investigation, Writing – review and editing. JJK: Conceptualization, Funding acquisition, Investigation, Supervision, Writing – original draft, and Writing – review and editing.

## Methods

### Cells, virus, and pharmacological reagents

HEK293T (293T, CRL-3216) and Vero (CCL-81) cells were obtained from American Type Culture Collection (ATCC) and maintained in Dulbecco’s modified Eagle’s medium (DMEM; 10-013-CV, Corning) supplemented with 10% heat-inactivated fetal bovine serum (FBS; 35-011-CV, Corning). Primary Dermal Fibroblast Normal, Human, Adult (HDFa) (PCS-201-012, ATCC) were maintained in Fibroblast Basal Medium (PCS-201-030, ATCC) with Fibroblast Growth Kit–Low Serum (PCS-201-041, ATCC). HAP-1 FASN WT and KO cells were obtained from Horizon Discovery and maintained in Iscove’s modified Dulbecco’s medium (IMDM; 15-016-CV, Corning) supplemented with 10% FBS. The following reagent was obtained through BEI Resources, NIAID, NIH: Cercopithecus aethiops kidney epithelial cells with high expression of human furin (Vero-Furin), NR-55312. Vero-Furin cells were maintained in DMEM with 10% FBS. All cells were grown in an incubator at 37 °C and 5% CO_2 a_nd maintained for less than 25 passages. The following reagent was obtained through BEI Resources, NIAID, NIH, as part of the WRCEVA program: Mayaro Virus, Guyane, NR-49911. The pharmacological inhibitors were obtained from sources listed in parenthesis and were resuspended in DMSO: Fasnall (SML1815, Sigma), TVB-2640 (35703, Cayman Chemicals), C75 (HY-123645, Medchem Express LLC), 2-bromopalmitate (2-BP; 21604, Millipore Sigma), A922500 (A129800, Aladdin Scientific), and ettomoxir (A482603, Ambeed). When working with 293T cells, tissue culture plates and microscopy slides were coated with 5μg/ml or 20μg/ml, respectively, of Poly-D-Lysine (P0899, Millipore Sigma).

### Viral propagation

Confluent Vero-Furin cells were infected at an MOI 0.01 with a low passage viral stock in Eagle’s minimum essential medium (EMEM, 30-2003, ATCC), 1X Earle’s Balanced salt solution (E7510, Sigma), and 2% FBS. After a 2-hour adsorption phase in the incubator, fresh medium was replenished, and cells were cultured until CPE is observed, approximately 48h.p.i. Cells were scraped and then media and cells were pulse vortexed for 30s prior to centrifugation at 1200xg for 10 minutes. Supernatant was filter sterilized with 0.45μm cellulose acetate (CA) syringe filter (76479-040, VWR) and aliquots were flash frozen on dry ice and stored at −80°C. Virus titers were determined via a plaque assay as described below.

### Viral infections

For pharmacological inhibition assays in 293T cells, 2×10^4 c^ells/well were seeded 24h prior to infection in 96-well plates. Cells were pre-treated with the designated inhibitors for 1h and then infected with MAYV at MOI 0.01 for 24h. Inhibitors were re-added to the media during infection. For pharmacological inhibition assays in HDFa cells, 5×10^4 c^ells/well were seeded 24h prior to infection in 96-well plates. HDFa cells were pre-treated with TVB-2640 for 1h and then infected with MAYV at MOI 1 for 1h. Inhibitors were re-added to the media during infection. Following viral adsorption, inoculum was removed, cells were washed once with 1xPBS, and fresh media with inhibitor was replenished for 24h. For other infections, 2.5×10^5 2^93T cells were seeded in 12-well plates 24h prior to infection at MOI 1, 5 or 10 for 1h. Following infection, the inoculum was removed, cells were washed once with 1xPBS and replenished with fresh media. Cells were incubated for 6h or 24h. For all experiments, cell culture supernatants were collected at the indicated time points to quantify virus titer via plaque assay.

### Plaque assay

Ten-fold dilutions of supernatants were prepared in DMEM with 2% FBS and 100 from each dilution was inoculated onto confluent Vero cell monolayers in 12-well plates. During the 1h viral adsorption period, an aqueous 0.6% UltraPure™ Agarose (16500500, Thermo Fisher Scientific) solution was prepared and dissolved by microwaving. The agarose solution was filter sterilized with a 0.22 μm CA syringe filter (76479-044, VWR) and cooled in a water bath at 56°C for 30 minutes. Subsequently, the agarose solution was mixed 1:1 in DMEM with 2% FBS and 1mL was added to each well. Following a 48h incubation, the cells were fixed with 10% paraformaldehyde (PFA) in 1xPBS for 1h. The agarose plugs were removed, and the cell monolayer was washed with 1xPBS and stained for 10 minutes with 0.05% crystal violet dissolved in 20% methanol. Finally, cells were rinsed with 1xPBS and plaques counted to determine the viral titer as PFU/mL. Fold-reduction was calculated by dividing the mean of the control group by that of the experimental group.

### CPE assay

HAP-1 FASN WT and KO cells were infected from MOI 1 to 0.008 (2 -fold dilutions) in 8 replicate columns for 24, 48 or 72h. An uninfected column was included as a control. After, cells were fixed with 4% PFA in 1xPBS for 1h, stained for 10 minutes with 0.05% crystal violet dissolved in 20% methanol, washed in 1xPBS and scored for CPE under the microscope. CPE was defined as cell detachment, rounding, and/or elongation across a well from a 96-well plate. The TCID_50_/mL was determined with the Reed-Muench method using the calculator from https://www.virosin.org/tcid50/TCID50.html (74).

### Sequence alignment and AlphaFold prediction modeling

The amino acid sequences for CHIKV nsP1 181/25 strain (UFI00979.1) and MAYV nsP1 Guyane strain (AAY45741.1) were obtained from NIH GenBank. The amino acid sequences had lengths of 535 aa for CHIKV and 536 aa for MAYV and were aligned with SIM in Expasy (SIB Swiss Institute of Bioinformatics)(75,76). AlphaFold3 (77) was used to predict the structures of nsP1 by including the S-palmitoyl modifications in residues 417, 418, and 419. Predictions were observed and aligned using PyMOL (The PyMOL Molecular Graphics System, Version 3.0.3 Schrödinger, LLC). Root mean square deviation (RMSD) values were obtained from aligned models and refined calculations after 5 cycles.

### Plasmids and infectious molecular clone (IMC) generation cloning

The pcDNA3.1-Flag-nsP1 construct had the nsP1 sequence of Mayaro virus strain MAYLC (DQ001069.1) (78) with an N-terminal start codon and Flag epitope tag (5’-ATGGACTACAAGGACGACGATGACAAG-3’), Kozak sequence immediately upstream (5’-GCCGCCACC-3’), stop codon downstream of nsP1 (5’-TAA-3’), and flanking 5’ EcoRI and 3’ XbaI restriction sites and adjacent regions from pcDNA3.1 (+). The WT and C417-419A constructs were synthesized as gene fragments (gBlocks, Integrated DNA Technologies), double-digested with EcoRI-HF (NEB, R3101) and XbaI (R0145, NEB) enzymes for ligation into pcDNA3.1(+) (V79020, Thermo Fisher Scientific) using T4 DNA ligase (M0202, NEB), according to manufacturer protocols. For single and double cysteine-to-alanine mutants, MAYLC Flag-nsP1 WT was amplified using flanking primers, ligated into pGEMT (A3600, Promega), and used as a template for single-primer site-directed mutagenesis as previously described (79). Mutant inserts were screened by Sanger sequencing (Genomics Shared Resources, OSUCCC-James), gel purified using the QIAquick gel extraction kit (28704, QIAGEN) and ligated into pcDNA3.1(+). Infectious molecular clones (IMCs) of MAYV strains TRVL 4675 and TRVL 15537 were kindly provided by Dr. James Weger-Lucarelli and are previously described (60). The TRVL 4675 IMC was chosen as a backbone to append an N-terminal Flag epitope tag to nsP1. Gene fragments (gBlocks, Integrated DNA Technologies) were synthesized containing the MAYV TRVL 4675 nsP1 sequence, wild-type or C417-419A, with an N-terminal start codon and Flag epitope tag (5’-ATGGACTACAAGGACGACGATGACAAG-3’), the native start codon removed, and 40 nucleotides of TRVL 4675 flanking nsP1. The Flag-nsP1 constructs were PCR amplified using Q5 High-Fidelity DNA Polymerase (M0491, NEB) and PCR purified (T1130S, NEB). Flag-nsP1 amplicons were then assembled with three overlapping fragments from the TRVL 4675 IMC using the NEBuilder HiFi Assembly Kit (E2621, NEB) following the manufacturers protocol. IMC assembly strategy was determined using NEBuilder Assembly Tool v2.10.8 set to an overlap of 25 nucleotides between fragments. pcDNA3.1-Flag-nsP1 constructs were transformed into NEB® 5-alpha Competent *E. coli* (C2988J, NEB) and IMC constructs were transformed into NEB® Stable Competent *E. coli* (C3040H, NEB) selected on LB-Agar with 50 μg/mL carbenicillin, cultured overnight in LB broth at 30°C with shaking, and plasmid DNA was extracted (12163, QIAGEN Plasmid Maxi Kit). Whole plasmid sequencing (Plasmidsaurus) was performed to verify sequence fidelity of all constructs prior to use. Gene fragments and primers used for plasmid construction, mutagenesis, and IMC assembly are provided in Supplementary Tables 1-3.

### Transfection, metabolic labeling, immunoprecipitations, and copper-catalyzed azide– alkyne cycloaddition

293T cells seeded at 8×10^5 c^ells/wells in 6-wells plates were reverse transfected with Lipofectamine 3000 (L3000008, Thermo Fisher Scientific) following the manufacturer’s protocol. For cells transfected with empty vector control (pcDNA3.1 +), pcDNA3.1-Flag-nsP1 WT or Flag-nsP1 C417-419A, expression occurred overnight prior to labeling. For cells transfected with TRVL 4675, Flag-nsP1 WT IMC or Flag-nsP1 C417-419A IMC, expression occurred for 48h prior to labeling. Cells were pre-treated with designated pharmacological inhibitors at the indicated concentrations for 6h in DMEM with 10% FBS. Afterward, media was removed, cells were washed with 1xPBS and labeled for 24h with 10 mM Alk-4, 50μM Alk-16, or DMSO in DMEM supplemented with 2% charcoal dextran-stripped FBS (35-072-CV, Corning) and 25 mM HEPES. Inhibitor was also added to the labeling media. For the remainder of the experiment, we followed the protocol by Yount et al (58). In summary, cells were washed in ice-cold 1xPBS and lysed in 1% Brij buffer (1% [w/v] Brij O10, 150 mM NaCl, 50 mM triethanolamine pH 7.4 with 1x EDTA-free complete protease inhibitor cocktail). Protein concentration was determined using the bicinchoninic acid (BCA) assay (A55861, Thermo Fisher Scientific). Flag precipitations used 450μg of cell lysate with anti-Flag Rabbit Polyclonal antibody (20543-1-AP, Proteintech) at 1:100 dilution and were mixed overnight at 4°C. After, 20 μl of washed protein G agarose resin (20398, Pierce) was added per sample and immunoprecipitation performed for 2h at RT while mixing. Antibody-bead complex was washed thrice with ice-cold lysis buffer, and complex was released by adding 22.5 μl of 4% SDS lysis buffer (150 mM NaCl, 50 mM triethanolamine, and 4% [w/v] SDS), and the click reaction was initiated by addition of 2.75 μl of click chemistry master mix composed of 0.5 μl of 5 mM 5-TAMRA Azide (CCT-1246-1, Vector Laboratories) in DMSO, 0.5 μl of 50 mM Tris-(2-Carboxyethyl)phosphine, Hydrochloride (TCEP; T2556, Thermo Fisher Scientific) in water, 0.5 μl of 50 mM CuSO4 (C1297, Sigma) in water, and 1.25 μl of 2 mM Tris ((1-benzyl-1H-1,2,3-triazol-4-yl)methyl)amine (TBTA; CCT-1061-100, Vector Laboratories) in 1:4 (v/v) DMSO/butanol. Reactions were incubated for 1h at RT and 10.8 μl of 4x Bolt^TM L^DS Sample Buffer (B0007, Thermo Fisher Scientific) with 5% beta-mercaptoethanol was added to samples. Samples were resolved in 10% Tris-glycine gels. To detect labeled proteins via fluorescence, the gel was destained in 40% distilled water, 50% methanol, 10% acetic acid (v/v) overnight at 4°C and visualized with the Azure 600 imaging system (Azure Biosystems).

### siRNA FASN knockdown and infection

ON-TARGET plus SMART pool of four siRNAs targeted against human *FASN* (L-003954-00-00010) and ON-TARGET plus non-targeting control siRNA (D-001810-01-05) were purchased from Horizon Dharmacon Reagents. 293T cells seeded at 2×10^5 c^ells/well were reverse transfected with 100nM FASN-targeting siRNA or 100nM NT siRNA using TransIT-X2® Dynamic Delivery System (MIR 6000, Mirus Bio) in 12-well plates. After 48h, cells were infected at MOI 0.01 and incubated for an additional 24h. Supernatants were collected for viral titer quantification via plaque assay. The cell monolayers were washed with 1xPBS and lysed on ice for 20 minutes using RIPA buffer supplemented with protease inhibitors. Cell lysates were cleared by centrifugation at 16,000xg for 16 minutes at 4°C and stored at −80°C for Western blot analysis.

### Western Blotting

Total protein concentrations were quantified using a BCA assay. For FASN expression analysis, 15 μg of total protein per samples was resolved on an 8% Tris-glycine gel. FASN and the actin loading control were analyzed on the same gel. For Flag-tag expression analysis, 10 μg of protein per sample was resolved on 10% Tris-glycine gels. Flag and the actin loading control were loaded on separate gels at identical protein concentrations and sample volumes. Gels were run for 1h with parameters set at 200V and 0.16A. Proteins were transferred onto 0.2 µm nitrocellulose membranes using the Trans-Blot Turbo RTA Mini Transfer Kit (1704270, BioRad) via the PowerBlotter Semi-Dry Transfer System (Thermo Fisher Scientific) according to the manufacturers pre-programmed settings. Membranes were blocked with 5% non-fat dry milk dissolved in 1xTris-buffered saline containing 0.1% Tween-20 (TBS-T) for 1h at RT. Anti-Flag tag monoclonal antibody (MA1-91878, Thermo Fisher Scientific) and anti-FASN (SAB4300700, Sigma) antibodies were used at 1:1000 and anti-actin (MA5-11869, Thermo Fisher Scientific) antibody was used at 1:2000. Primary antibodies were incubated overnight at 4°C while rocking. The secondary antibodies StarBright Blue 520 Goat Anti-Rabbit IgG (2005869, BioRad) and StarBright Blue 700 Goat Anti-Mouse IgG (2004158, BioRad) were used at a concentration of 1:2000 and incubated for 1h. Western blots were visualized using the Azure 600 Imaging System (Azure Biosystems). Densitometry quantification of protein bands and conversion of fluorescent images to grayscale was performed using Fiji software (80).

### RT-qPCR

Total RNA from infected cells was extracted using the Direct-zol RNA Miniprep Kit (R2051, Zymo Research). RT-qPCR was performed with Luna® Universal One-Step RT-qPCR Kit (E3005X, New England Biolabs) according to manufacturer’s instructions and thermocycling conditions. MAYV expression was quantified with the following primers: forward, 5′-TCAGCCGCCCTTTCCTTCAT-3′; reverse 5′-TTGCGCCCAAGTCATTGCAG-3′. The *18S rRNA* subunit was amplified as an endogenous reference control for sample normalization using the following primers: forward, 5′-CAGCCACCCGAGATTGAGCA-3′; reverse, 5′-TAGTAGCGACGGGCGGTGTG-3′. Thermocycling was carried out for 40 cycles, with annealing temperatures of 57°C and 58°C for MAYV and *18S rRNA*, respectively. Relative gene expression levels were calculated using the comparative Ct method (ΔΔCt) and MOI 10 was set as the reference value.

### Microscopy

293T cells seeded at 7×10^4 c^ells/well in 8-well chamber slides (PEZGS0896, Sigma) 24h prior to infection and cells were infected at MOI 1 for 24h. Afterwards, cells were fixed in 4% PFA for 10 minutes at RT, washed in 1xPBS, permeabilized with 1xPBS + 0.1% Triton X-100 for 20 minutes, and blocked with 3% BSA in 1xPBS for 1h. For intracellular staining, cells were incubated with anti-FASN (ab128870, Abcam) monoclonal antibody used at 1:500 and anti-dsRNA (J2; Kf-Ab01299-2.0, Kerafast) used at 1:60 overnight in 4°C while rocking. Cells were washed with 1xPBS + 0.1% Tween-20 (PBS-T), incubated with F(ab’)2-Goat anti-Mouse IgG (H+L) Cross-Adsorbed Secondary Antibody, Alexa Fluor™ 488 (A-11017, Thermo Fisher Scientific) and Goat anti-Rabbit IgG (H+L) Cross-Adsorbed Secondary Antibody, Alexa Fluor™ 594 (A-11012, Thermo Fisher Scientific) at 1:1000 for 1hr, washed with 1xPBS-T and nuclei stained with 150nM of DAPI for 15 minutes. Lastly, the cells were washed with 1xPBS-T and mounted with Aqua-Poly/Mount (18606, Polysciences). All washing steps were done three times for 5 minutes. Cells were imaged using a 40x objective lens with a Leica DM6 B upright fluorescent microscope. Brightness and contrast was adjusted uniformly in DAPI channels using Fiji software (80) to enhance nucleus visibility. Nonlinear adjustments were not performed on dsRNA or FASN channels.

### Cell viability assays

293T cells seeded at 1.5×10^4 c^ells/well or HDFa cells seeded at 5×10^4 c^ells/well 24h prior were exposed to inhibitors in 2-fold dilutions (100 μM-0.3 μM) in triplicates for 24h. Vehicle (DMSO) control wells were included. Cell viability was measured using the Calcein AM assay (C3100MP, Thermo Fisher Scientific) following the manufacturer’s protocol and reading conditions included well scan at excitation 495nm and emission 530nm, pattern: fill, points per well: 9, density: 3, point spacing of approx. 1.51, bottom read, PMT gain: high, Flashes per read: 10. All plates were read with a SpectraMax i3x Multi-Mode Microplate Reader (Molecular Devices).

### Statistics and figure generation

Statistically significant differences were determined using the Mann-Whitney test in GraphPad Prism 11 (GraphPad Software Inc., La Jolla, CA). Adobe Illustrator was used for figure organization. Schematics made with BioRender. (Loperena, P. (2026) https://BioRender.com/gpa0vuk.)

